# T cells expanded from renal cell carcinoma are tumor-reactive but fail to produce IFN-γ, TNF-α or IL-2

**DOI:** 10.1101/2020.05.17.098558

**Authors:** Saskia D. van Asten, Rosa de Groot, Marleen M. van Loenen, Jeroen de Jong, Kim Monkhorst, John B.A.G. Haanen, Derk Amsen, Axel Bex, Robbert M. Spaapen, Monika C. Wolkers

**Affiliations:** Sanquin Research, Dept. of Immunopathology, Amsterdam, The Netherlands; Landsteiner Laboratory, Amsterdam UMC, University of Amsterdam, Amsterdam, The Netherlands; Sanquin Research, Dept. of Hematopoiesis, Amsterdam, The Netherlands; Oncode Institute, Utrecht, The Netherlands; The Netherlands Cancer Institute-Antoni van Leeuwenhoek Hospital (NKI-AvL), Dept. of Pathology, Amsterdam, The Netherlands; NKI-AvL, Dept. of Medical Oncology, Amsterdam, The Netherlands; NKI-AvL, Dept. of Urology, Amsterdam, The Netherlands; Royal Free London NHS Foundation Trust, UCL Division of Surgery and Interventional Science, London, United Kingdom

**Keywords:** TIL therapy, RCC, immunotherapy, immune composition, dysfunctional T cells, CD137

## Abstract

Metastatic renal cell carcinoma (RCC) has a poor prognosis. Recent advances have shown beneficial responses to immune checkpoint inhibitors, such as anti-PD-1 or anti-PD-L1 antibodies. As only a subset of RCC patients respond, alternative strategies should be explored. Patients refractory to anti-PD-1 therapy may benefit from autologous tumor infiltrating lymphocyte (TIL) therapy. Even though efficient TIL expansion was reported from RCC lesions, it is not well established how many RCC TIL products are tumor-reactive, how well they produce pro-inflammatory cytokines in response to autologous tumors, and whether their response correlates with the presence of specific immune cells in the tumor lesions.

We here compared the immune infiltrate composition of RCC lesions with that of autologous kidney tissue of 18 RCC patients. T cell infiltrates were increased in the tumor lesions, and CD8^+^ T cell infiltrates were primarily of effector memory phenotype. Nine out of 16 (56%) tested TIL products we generated were tumor-reactive, as defined by CD137 upregulation after exposure to autologous tumor digest. Tumor reactivity was found in particular in TIL products originating from tumors with a high percentage of infiltrated T cells compared to autologous kidney, and coincided with increased *ex vivo* CD25 expression on CD8^+^ T cells. Importantly, although TIL products had the capacity to produce the key effector cytokines IFN-γ, TNF-α or IL-2, they failed to do so in response to autologous tumor digests. In conclusion, TIL products from RCC lesions contain tumor-reactive T cells. Their lack of tumor-specific cytokine production requires further investigation of immunosuppressive factors in RCC and subsequent optimization of RCC-derived TIL culture conditions.

## Introduction

Patients with metastatic renal cell carcinoma (RCC) have a poor 5-year survival rate.^1^ Several lines of evidence suggest that RCC patients may benefit from immunotherapy.^1,2^ In the past, treatment with high dose IL-2 resulted in partial responses. However, severe acute toxicities were common.^3–5^ Allogeneic stem cell transplantation also elicited complete responses in several RCC patients,^6,7^ a finding that could not be reproduced by others.^8–10^ Whilst the degree of T cell infiltration in RCC inversely correlated with the patient’s survival, recent advances with checkpoint inhibitors to reinvigorate T cell responses yielded robust response rates in RCC patients.^11,12^ Antibodies directed against PD-1 or its ligands PD-L1 and PD-L2 in combination with anti-CTLA-4 or axitinib, a VEGFR tyrosine kinase inhibitor, resulted in 50% objective response rates of metastatic clear-cell RCC patients, and led to complete responses in 10% of the patients.^13,14^ This combination therapy has therefore replaced anti-VEGF targeted therapy as the new standard of care for treatment of naïve RCC patients.

Another emerging immunotherapeutic treatment is the administration of *ex vivo* reprogrammed tumor infiltrating lymphocytes (TILs). This adoptive TIL therapy induced objective responses in almost half of treated metastatic melanoma patients, and complete responses in 15-20% of patients.^15–17^ Furthermore, TIL infusion yielded an objective response rate of 38% in melanoma patients that were refractory to anti-PD-1 therapy, indicating that TIL therapy represents an alternative treatment for patients who fail to respond to immune checkpoint blockade.^18^ The success rate of TIL therapy to treat metastatic melanoma has sparked the interest to develop TIL therapy for other solid tumors, such as ovarian, colon, liver, breast and non-small cell lung cancers.^19–24^ TIL products have also been generated from RCC lesions.^25–28^ Although the expansion rate of RCC-derived TILs was comparable to that of melanoma-derived TILs, the *in vitro* response rates to autologous tumors were highly variable,^25–28^ a characteristic that is not well understood. It would be useful to be able to predict from the composition of the initial tumor infiltrate which TIL products will be tumor-reactive. However biomarkers that allow such prediction have not been identified. Furthermore, it is unclear whether those RCC-derived TIL products that respond to tumors contain T cells that co-produce multiple effector molecules in response to autologous tumors. Such polyfunctionality is considered a prerequisite for successful anti-tumor T cell responses as well as for TIL therapy.^29,30^

In this study, we characterized the patient-specific RCC immune cell composition compared to autologous non-tumor kidney tissue. Irrespective of high inter-patient variation, we found that T cells, especially of the effector memory subtype, constituted the major immune cell type in the RCC tumors, that were predominantly of the clear cell subtype. RCC-derived T cells responded to the autologous tumor digest by CD137 upregulation in 9 out of 16 (56%) patients. Higher frequencies of tumor-reactive T cells were found in TIL products generated from tumor lesions with high T cell infiltrates, in particular when CD8^+^ T cells expressed high levels of CD25. Strikingly, even though expanded TILs had the capacity to produce all key cytokines upon PMA/ionomycin stimulation, they failed to do so in response to autologous tumor tissue.

## Methods

### Materials and solutions

Several solutions were used in sample processing. Collection medium consisted of 50µg/ml gentamycin (Sigma-Aldrich), 2% Penicillin-Streptomycin (P/S), 12.5µg/ml Fungizone (Amphotericin B, Gibco) and 20% fetal calf serum (FCS) (Bodego). Digestion medium consisted of 30 IU/ml collagenase IV (Worthington), 1% FCS and 12.5µg/ml DNAse (Roche) in IMDM (Gibco). Washing medium consisted of 2% FCS and 2% P/S in RPMI 1640 (Gibco). FACS buffer contained 2% FCS and 2mM EDTA in PBS. Red blood cell lysis buffer consisted of 155 mM NH4Cl, 10 mM KHCO3 and 0.1 mM EDTA (pH 7.4) in PBS. T cell culture medium consisted of 5% human serum (Sanquin), 5% FCS, 50 µg/ml gentamycin and 1.25 µg/ml fungizone in 20/80 T cell mixed media (Miltenyi Biotech). Freezing medium consisted of 10% DMSO (Corning) and 30% FCS in IMDM.

### Sampling of tumor and non-tumor kidney tissue

Tumor and spatially distant non-tumor kidney tissue were collected from 20 RCC patients undergoing a nephrectomy from April 2016 to March 2018. Patient 08 was excluded from this study as the collected tumor piece proved too small for isolation of sufficient cell numbers for *ex vivo* analysis. A patient with oncocytoma (patient 11) was excluded from all analysis because we aimed to analyze malignant material only. From patient 05 we could only collect tumor tissue. This patient was therefore excluded from non-tumor and tumor paired comparisons. The patient characteristics of the 18 included patients are shown in Table 1. The cohort included 5 female and 13 male patients. 14 Tumors were classified as clear cell carcinoma, three as chromophobe and one as not otherwise specified (NOS). The tumor diameters ranged from 5.2 cm to 16.5 cm. The patient age at time of surgery ranged from 51 to 79, with a mean age of 63. Seven patients were smokers. The study was performed according to the Declaration of Helsinki (seventh revision, 2013), and executed with approval of the Institutional Review Board of the Dutch Cancer Institute (NKI), Amsterdam, the Netherlands.

**Table 1.** patient and tumor characteristics.

| Patient | Sex (M/F) | Age at surgery | Smoker | Pretreatment | RCC type | Fuhrman grade | Max tumor diameter (cm) | # tumors within kidney | PD-L1 TC (%) | PD-L1 IC (IC0-3) |
| --- | --- | --- | --- | --- | --- | --- | --- | --- | --- | --- |
| 01 | M | 61 | yes | sunitinib | ccRCC | 3 | 9 | 1 | 0 | IC1 |
| 02 | M | 59 |  |  | chRCC | NA | 13 | 2 | 60 | IC1 |
| 03 | M | 66 |  | sunitinib | ccRCC | 2 | 9 | 1 | 0 | IC2 |
| 04 | M | 75 |  | sunitinib | ccRCC, metastasis adrenal gland | 2 | 8.5 | 1 | ND | ND |
| 05 | F | 71 |  |  | ccRCC | 2 | 8.5 | 2 | 0 | IC0 |
| 06 | M | 59 |  |  | ccRCC | 3 | 8 | 2 | 0 | IC0 |
| 07 | M | 55 | yes |  | NOS, metastasis adrenal gland | NA | 13 | 1 | 1 | IC0 |
| 09 | M | 54 | yes | axitinib + avelumab | ccRCC, metastatic | 3 | 7 | 1 | 90 | IC2 |
| 10 | F | 61 |  |  | ccRCC | 1 | 14 | 1 | 0 | IC0 |
| 12 | F | 78 |  |  | ccRCC | 4 | 5.5 | 1 | 10 | IC0 |
| 13 | M | 80 |  |  | chRCC | NA | 7 | 1 | 0 | IC0 |
| 14 | M | 57 |  |  | ccRCC | 2 | 4.5 | 1 | 1 | IC2 |
| 15 | M | 61 | yes |  | ccRCC | 3 | 9.2 | 1 | 0 | IC2 |
| 16 | M | 57 |  |  | ccRCC | 3 | 8.5 | 1 | 0 | IC1 |
| 17 | M | 63 |  | pazopanib | chRCC metastatic | 4 | 16.5 | 1 | 0 | IC2 |
| 18 | F | 64 |  | sunitinib | ccRCC | 3 | 9.5 | 1 | 0 | IC1 |
| 19 | F | 65 | yes |  | chRCC | 2 | 8.5 | 1 | 30 | IC0 |
| 20 | M | 52 | yes | sunitinib | ccRCC, metastatic | 1 | 7 | 1 | 0 | IC2 |

Abbreviations: M = male, F = female, ccRCC = clear cell RCC, chRCC = chromophobe RCC, NOS = not otherwise specified, NA = not applicable, ND = not determined, TC = tumor cell, IC = immune cell, IC0-3 = PD-L1 immune cell score 0-3

The tissue was obtained directly after surgery, stored in collection medium, weighed, and completely processed within 5 hours, except for patients 12 and 13, which were processed within 18 hours.

### Isolation & digestion

We used our standardized isolation, digestion and TIL culture procedures as described before^21^. In short, the tissues were chopped into small pieces of ∼1mm^3^ and incubated for 45 minutes at 37 °C in 5 ml digestion medium/g tissue. After digestion, cell suspensions were centrifuged for 10 min at 360 g. The pellets were washed with washing medium. After removal of the washing medium by centrifugation, cell pellets were vigorously suspended in FACS buffer to loosen the cells from the tissue. The cell suspensions were filtered over an autoclaved tea filter, followed by filtering over a 100 µm cell strainer (Falcon). The pelleted flow-through was treated with Red blood cell lysis buffer for 15 minutes at 4°C. The cell suspensions were again washed with FACS buffer before counting. The cells were manually counted in trypan blue solution (Sigma). 1-2 million live cells were used per flow cytometry panel (2-4 million total), 1 million live cells for TIL cultures, and the remaining cells were cryo preserved.

### Ex vivo flow cytometry

Cells were stained at 4°C in FACS buffer for 30 minutes in U-bottom 96-well plates at 1-2 million cells per panel. For the immune cell subset analysis, the staining mix included the following antibodies: CD11c PerCP-Cy5.5, CD15 BV605, CD3 BV510, CD11B BV421, CD274 PE (all Biolegend), anti-HLA-DR FITC, CD45 BUV805, CD19 BUV737, CD20 BUV737, CD16 BUV496, CD1a BUV395, CD33 AF700 (all BD Biosciences), CD14 PE-Cy7 (eBioscience) and LIVE/DEAD™ Fixable Near-IR Stain (Invitrogen). For phenotypic analysis of T cells, the staining mix included the following antibodies: CD3 PerCP-Cy5.5, PD-1 FITC, CD56 BV605, CD27 BV510, CCR7 BV421, CD103 PE-Cy7, CD25 PE (all Biolegend), CD8 BUV805, CD45RA BUV737, CD4 BUV496, CD69 BUV395 (all BD Biosciences) and LIVE/DEAD™ Fixable Near-IR Stain. After antibody incubation at RT degrees for 20 min, cells were centrifuged for 4 min at 350 g. Cell pellets were washed twice with FACS buffer. Cells were fixed using the fixation solution of the anti-human FoxP3 staining Kit (BD Biosciences). After fixation for 30 minutes, cells were washed twice with FACS buffer, and then once with the permeabilization buffer of the anti-human FoxP3 staining kit. For immune cell subset analysis, cells were stained with anti-FOXP3 APC and for the T cell phenotype analysis with anti-CD68 APC in permeabilization buffer for 30 minutes. Cell suspensions were washed twice with permeabilization buffer, taken up in FACS buffer and filtered using Filcon Syringe-Type 50 µm filters (BD biosciences) right before measurement. Samples from patients 01–09 were measured on a LSRFortessa flow cytometer and samples from patients 10–20 on a FACSymphony (both BD Biosciences).

### Immunohistochemistry

Formalin-fixed, paraffin-embedded tumor tissue samples were stained for PD-L1 (clone 22C3, Agilent) using the Ventana Benchmark Ultra. Membranous PD-L1 expression of any intensity on viable tumor cells was scored as a percentage of at least 100 tumor cells. The percentage of PD-L1 positive immune cells was scored as the proportion of tumor area that is occupied by PD-L1 staining of immune cells of any intensity and categorized as follows: 0% = IC0, 0% > IC1 < 5% > IC2 < 10% < IC3.

### TIL culture

Freshly isolated live cells were cultured in 24 well plates (10^6^ cells per well) in T cell culture medium supplemented with 6000 IU/ml IL-2 for ∼2 weeks at 37 °C 5% CO_2_. Medium was refreshed approximately every 4 days. Wells were split when a monolayer of cells was observed in the entire well. After this initial culture period, cells were cultured using a standard rapid expansion protocol (REP) for another two weeks: the culture medium was replaced with T cell culture medium supplemented with 3000 IU/ml IL-2 and anti-CD3 (clone OKT3; Miltenyi Biotech), also containing 8-10 × 10^6^ irradiated (40 Gray) PBMCs pooled from 10 healthy donors. Cells were passaged when multiple clusters of dividing T cells were observed, which was usually once or twice during the two-week REP culture period. The expanded T cells were cryo-preserved in freezing medium.

### Tumor reactivity assay

Expanded T cells were thawed in RPMI containing 2% FCS at 37°C. Cells were washed twice with T cell culture medium and centrifugation at 350 g for 5 min. The number of living trypan blue negative cells was determined using a hemocytometer. Cells were pre-stained in FACS buffer containing CD8 BUV805, CD4 BUV496 and CD3 BUV395 (clone SK7, BD biosciences) for 30 min at 4°C, washed twice and taken up in T cell culture medium. 100,000 expanded live cells were assayed in T cell culture medium alone (= medium control) or co-cultured with either thawed tumor or kidney digest containing 50,000 to 100,000 living cells or exposed to PMA/ionomycin (10 ng/ml PMA, 1 µg/ml ionomycin). After one hour, Brefeldin A (1:1000, Invitrogen) and 1:1000 monensin (eBioscience) were added. After six hours, cells were washed with FACS buffer before staining with anti-PD-1 BV421 (BD biosciences), CD154 BV510 (Biolegend) and LIVE/DEAD™ Fixable Near-IR Stain for 30 minutes followed by centrifugation for 4 min at 350 g. Cell pellets were washed twice with FACS buffer. Cells were fixed using fixation solution (BD, FoxP3 staining kit) for 30 min and washed twice using FACS buffer. Cells were kept overnight at 4°C, washed with permeabilization buffer and stained with anti-TNF-α AF488, CD137 PE-Cy7, anti-IFN-γ PE (all Biolegend) and anti-IL-2 APC (Invitrogen) in permeabilization buffer for 30 min. Cells were washed twice with permeabilization buffer and analyzed in FACS buffer on a BD FACSymphony.

### Analysis

Flowjo version 10.1 (Tree Star) was used to analyze flow cytometry data. Further analysis of the percentages of the stated phenotypes, data transformations, generation of figures and statistical calculations were all performed in R. *P*-values were determined by indicated statistical tests and depicted using the following symbols: p<0.05 = *, p<0.01 =**, p<0.001 = ***, n.s. = not significant.

## Results

### Clear cell RCC lesions are highly infiltrated by T cells

To determine the composition of immune cells infiltrating RCC lesions, we collected both tumor and non-tumor sections from resected kidneys of 18 RCC patients, ranging from Fuhrman grade 1 to 4 and with varying PD-L1 expression on tumor and immune cells (IC) (Table 1). Roughly half of the patients had received pretreatment with protein tyrosine kinase inhibitors, while the other half were treatment naive (Table 1). The non-tumor section (hereafter referred to as kidney tissue) was dissected as spatially distant as possible from the tumor. Single cell suspensions were generated from the collected tissue by physical disruption and digestion with DNAase and collagenase IV. After extensive filtration, the number of live cells was determined. The yield of cells per mg tissue derived from tumor lesions and kidney tissue was similar (Fig. 1A).

**Fig 1.**
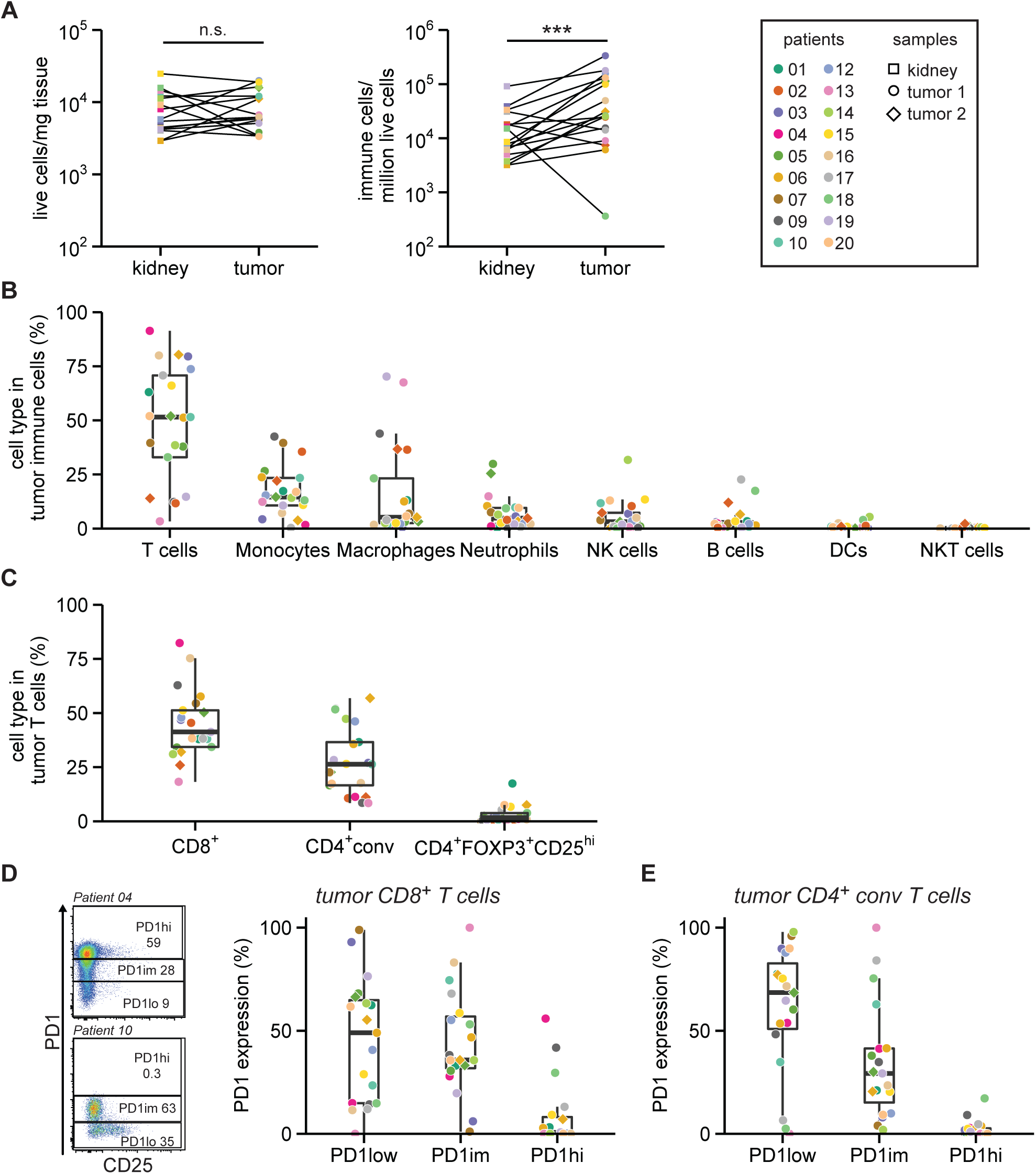
T cells are enriched in RCC lesions. **A**. Tissue from RCC lesions and distal non-tumor kidney tissue was collected from 18 RCC patients. The number of living cells in single cell suspensions were determined using trypan blue (left panel). The number of T cells, monocytes, macrophages, neutrophils, natural killer (NK) cells, B cells, dendritic cells (DCs) and NKT cells were determined by flow cytometry. Collectively these cell types are defined as immune cells (middle panel). Statistical significance was determined by Wilcoxon signed-rank test. Right panel: Patient color coding and sample shape coding, which is used throughout all figures. **B**. Percentage of indicated immune cell subsets of the total immune cells in tumor digests. **C**. T cell subsets as a percentage of total CD3^+^ T cells in the tumor digest as determined by flow cytometry. CD4^+^ conventional (conv) T cells are defined as non-FOXP3^+^CD25^hi^ (see Fig. S3). **D**. Gating strategy (left panel) of two representative patients to define high (hi), intermediate (im) and low (lo) PD-1 expression in tumor-infiltrating CD8^+^ T cells and summary (right panel). **E**. Similar gating was used for CD4^+^ conv T cells (E).

We next quantified the immune infiltrates by flow cytometry. We measured T cells, monocytes, macrophages, neutrophils, NK cells, B cells, dendritic cells (DCs) and NKT cells (gating in Fig. S1). Overall, higher numbers of immune cells were found in tumor lesions compared to paired kidney tissue (Fig. 1A). Similar to the kidney tissue, T cells constituted the major immune cell subset in RCC lesions, with an average of 48 ± 27% of the immune infiltrate (Fig. 1B; Fig. S2A). Also, monocytes and macrophages were >20% present in the infiltrate in tumor tissue of 6 and 5 patients respectively (Fig. 1B). The tumor infiltrates contained on average less than 5% of neutrophils, NK cells, B cells, DCs and NKT cells (Fig. 1B). Interestingly, tumors of the four patients with the lowest percentages of T cells (patient 02, 09, 13, and 19) also contained the highest percentages of macrophages (Fig. S2B). Of note, three of these patients were diagnosed with chromophobe RCC (Fig. S2B).

The high proportion of T cells in the tumor lesions led us to further explore these immune cell subsets (gating in Fig. S3A). Overall, CD8^+^ T cells were more abundant in tumor lesions than CD4^+^ conventional (conv) T cells (Fig. 1C). FOXP3^+^ CD25^hi^ T cells contributed on average 2.4 ± 3.2% to the total T cell population in RCC lesions (Fig 1C). These percentages of FOXP3^+^ T cells are substantially lower than those measured in melanoma or ovarian cancer lesions (∼15% of CD3^+^ T cells).^21^

High expression levels of PD-1 (PD-1hi) on CD8^+^ T cells in solid tumors is reportedly often indicative for tumor reactivity.^31^ Most RCC infiltrating CD8^+^ and CD4^+^ conv T cells showed low (PD-1low) or intermediate PD-1 (PD-1im) expression (Fig. 1D, E). Nevertheless, 6 tumors contained more than 5% PD-1hi CD8^+^ T cells, which in 2 cases (patients 09 and 18) coincided with the presence of PD-1hi CD4^+^ conv T cells. In conclusion, high percentages of CD8^+^ and CD4^+^ conv T cells with variable PD-1 expression levels are present in clear cell RCC lesions.

### RCC infiltrating T cells display an effector memory phenotype

We next determined the differentiation and activation status of tumor infiltrating T cells compared to T cells from paired kidney tissue. We analyzed the expression of CD45RA, CD27, CCR7, CD103 and CD69 to differentiate between naïve, effector and memory subtypes, and of CD25 and PD-1 to define T cell activation and exhaustion (gating in Fig. S3B). To prevent the inclusion of unreliable data, all populations included in the PCA contained at least 200 cells. We visualized the relationship between different patient samples and within patients by applying principal component analysis (PCA) on the percentage of T cell subtypes and the percentage of activation/exhaustion marker expression within these T cell subsets (Fig. 2A and B). Distances between T cells from tumor and kidney samples of individual patients seemed limited (Fig. 2A), suggesting that TILs resemble T cells from the corresponding kidney more than TILs from other patients. Nonetheless, PC2 potently separated T cells from tumor and kidney sample groups (Fig. 2B). We therefore examined which variables contributed most to PC2. Samples with a high PC2 score were enriched for CD27^-^CD45RA^-^ and CCR7^-^CD45RA^-^ expressing CD8^+^ and CD4^+^ conv effector memory T cells (T_EM_) compared to samples with a low PC2 score, which contained more naïve (CD27^+^CD45RA^+^ and CCR7^+^CD45RA^+^) expressing CD8^+^ and CD4^+^ conv T cells (Fig. 2C). Specific analyses of these parameters showed that both CD8^+^ and CD4^+^ conv T cells in tumor tissue were enriched for the T_EM_ phenotype (CD27^-^CD45RA^-^) compared to kidney T cells (Fig. 2D and E). Also, naïve CD8^+^ T cells (CD27^+^CD45RA^+^) were significantly reduced in tumors compared to CD8^+^ T cells in the kidney (Fig. 2D). In addition, samples with a low PC2 score were enriched for CD25 expressing CD8^+^ and CD4^+^ conv T cells (Fig. 2C). Indeed, tumor-derived CD8^+^ T cells and CD4^+^ conv T cells (which predominantly exhibit relatively high PC2 scores) had lower percentages of CD25 expression than T cells present in the kidney tissue (Fig. 2F and G). Although high CD25 expression is a hallmark of FOXP3^+^ Tregs, the overall percentage of infiltrated FOXP3^+^ CD25hi T cells did not significantly differ between tumor and kidney tissue (Fig. 2H).^32^ In conclusion, tumor-infiltrating T cells are enriched for a T_EM_ phenotype but have lower CD25 levels compared to T cells isolated from paired kidney tissue.

**Fig 2.**
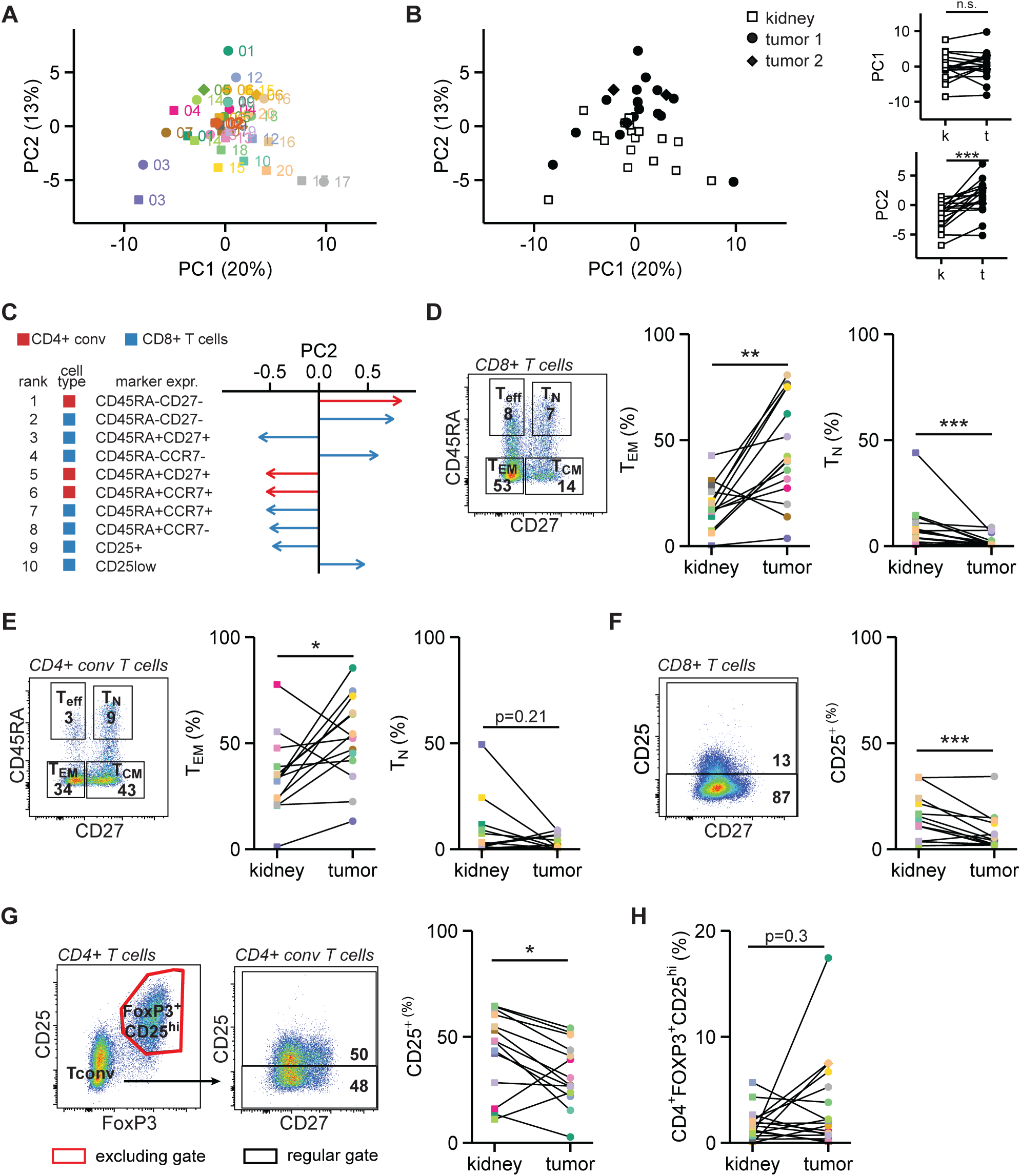
RCC-derived T cells are phenotypically different from kidney T cells. **A-B**. The activation and memory profiles of CD8^+^, CD4^+^ conv T cells and CD4^+^FOXP3^+^CD25^hi^ cells were determined by flow cytometry. These profiles included three PD-1 levels (low, intermediate, high), CD25 expression, CD103^+^CCR7^-^, CD103^-^CCR7^-^, and all combinations of CD27/CD45RA, CCR7/CD45RA and CD103/CD69 (see Fig. S3). Dimensions of these data were reduced using principal component analysis (PCA). The first two principal components (PC) are shown as colored by sample type (**A**) and by patient (**B**). **C**. The ten phenotypes that contributed most to variance in PC2. Colors indicate whether a phenotype was observed in CD4^+^ conv T cells (red) or CD8^+^ T cells (blue). **D**. Left panel: gating for memory phenotypes in CD8^+^ T cells. Middle and right panels: effector memory and naïve T cell percentages within CD8^+^ T cells. **E**. As in **D**, but then for CD4^+^ conv T cells. **F**. CD25 expression gating (left panel) and quantification (right panel) for CD8^+^ T cells. **G**. As in **F**, for CD4^+^ conv T cells. **H**. Percentage of CD4^+^FOXP3^+^CD25^hi^ cells in CD3^+^ T cells. For **D-H** only paired samples were plotted. Statistical significance in panels **A** and **D-H** was determined by Wilcoxon signed-rank test.

### Effective expansion of functionally active RCC TILs

To determine the expansion capacity of RCC TILs, we cultured tumor digests with 6000 IU/ml IL-2 for 14 days, followed by a 14 day rapid expansion protocol (REP)^33^ using soluble anti-CD3 antibodies and 3000 IU/ml IL-2. For comparison, we also expanded T cells isolated from kidney tissue. The expansion rates were similar between kidney-and tumor-derived T cells, suggesting that immunomodulatory factors that were potentially present in the tumor digest did not substantially affect the T cell proliferation potential in vitro (Fig. 3A). On average, the ratio of CD4^+^ to CD8^+^ T cells did not alter significantly upon T cell expansion (Fig. 3B; gating in Fig. S4A). Nonetheless, this ratio was altered in individual patients during culture (Fig. S4B).

**Fig 3.**
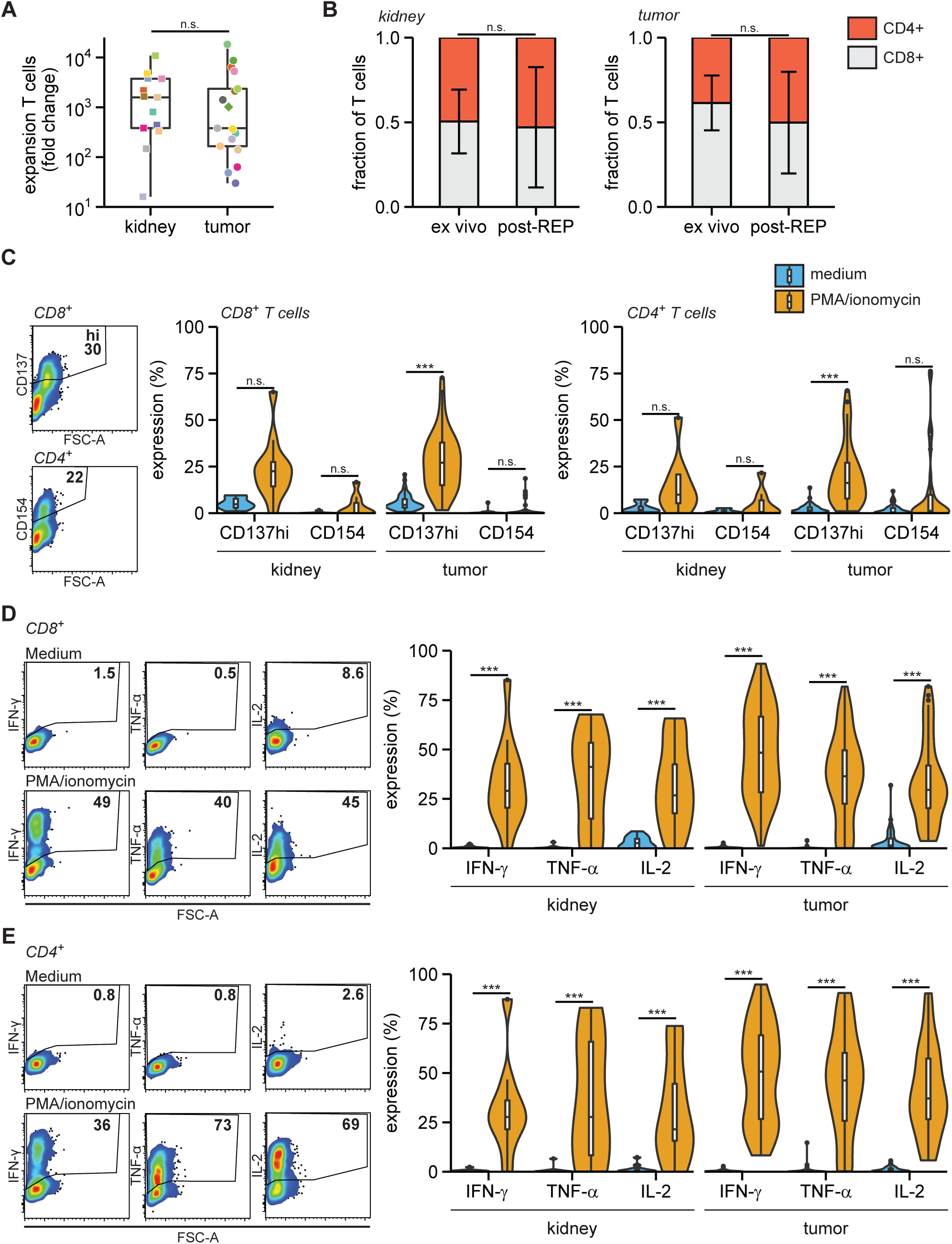
Expanded RCC TILs are capable cytokine producers. Tumor and kidney digests were cultured for two weeks in 6000 IU/ml IL-2, followed by two weeks of REP. **A**. Fold change of number of T cells post-REP compared to *ex vivo*. **B**. Fraction of CD8^+^ and CD4^+^ T cells from kidney (left panel) and RCC (right panel) *ex vivo* and post-REP (individual datapoints in Fig. S4B). **C-E**. Post-REP TILs were re-stimulated with PMA/ionomycin for 6 h and analyzed for expression of intracellular CD137 and surface CD154 (**C**), expression of intracellular cytokines in CD8+ T cells (**D**) and in CD4+ T cells in (**E**). Statistical significance was determined by Wilcoxon signed-rank test for panels **A** and **B** and by ANOVA with Tukey’s post hoc test for panels **C-E**.

Next, we measured the activation potential of the expanded TIL products. We stimulated TILs with PMA/ionomycin for 6h and measured the expression of the activation markers CD137/4-1BB, and of CD154/CD40L, two markers that indicate T cell receptor-dependent activation.^34–36^ In particular, the expression of CD137 was substantially induced in expanded CD8^+^ and CD4^+^ T cells generated from both kidney and tumor tissue (Fig. 3C; gating in Fig. S4A and C). Furthermore, the key pro-inflammatory cytokines IFN-γ, TNF-α and IL-2 were effectively produced by both kidney and tumor tissue-derived CD8^+^ and CD4^+^ T cells (Fig. 3D and E; gating in Fig. S4D). In conclusion, TILs from RCC lesions could be effectively expanded and were generally able to produce inflammatory cytokines in response to pharmacological stimulation.

### Expanded TILs are tumor-reactive

We next tested the tumor reactivity of expanded TILs from 16 RCC lesions and -when available-from kidney tissue. We co-cultured the TIL products with autologous tumor digests for 6h and measured the induction of CD137 and CD154. To define the base line CD137 and CD154 expression, we cultured the expanded TILs in medium alone and exposed them to autologous kidney digests. Induction of CD154 expression on CD4^+^ T cells was scarce in any of the culture conditions (Fig. 4A and S4E). Conversely, CD137 expression was significantly increased upon co-culture with tumor digest, in particular in CD8^+^ T cells (Fig. 4A and S4F). In fact, in 9 out of 15 analyzed individual patients (60%), the CD137 expression on TILs was higher upon coculture with tumor tissue digest than with the autologous kidney tissue digest (Fig. 4B; kidney digest was lacking for patient 05). This finding indicates that RCC TIL products specifically responded to tumor tissue digests.

**Fig 4.**
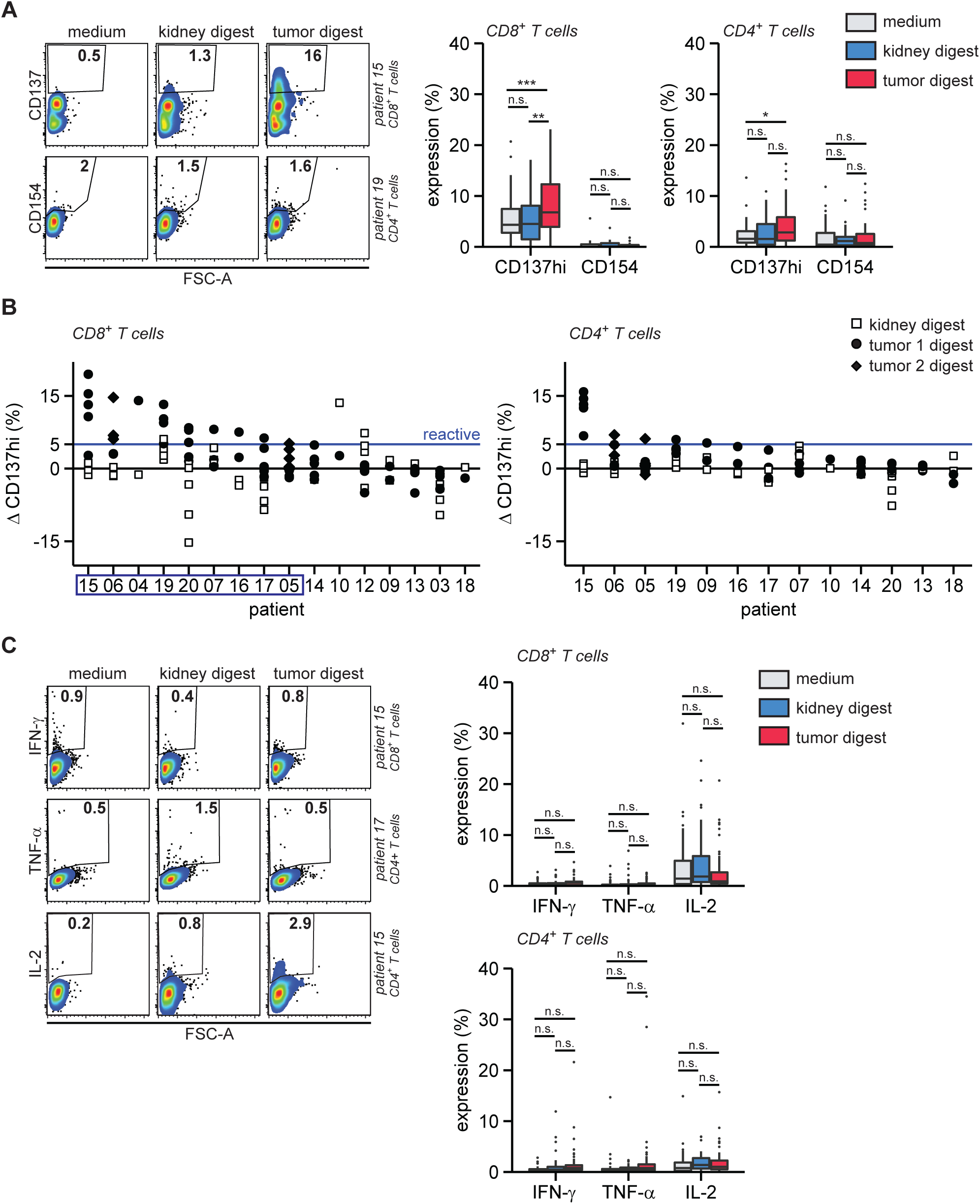
RCC TILs are tumor-reactive but do not produce key cytokines. Post-REP TILs were first stained for CD3, CD4 and CD8, then cultured in medium with or without autologous kidney or autologous tumor digests and analyzed by flow cytometry for CD137, CD154, IFN-γ, TNF-α or IL-2 after 6h of coculture. **A**. Intracellular CD137 and surface CD154 expression, representative examples (left panels) and summary of quantifications (right panels). **B**. The difference (Δ, Delta) in high CD137 (CD137hi) expression between co-culture of TILs with tissue digest and medium control (tissue digest – medium). TIL products with ΔCD137hi > 5% (blue line) in CD8^+^ T cells against tumor digests were considered tumor-reactive. Patients from which tumor-reactive TIL products were produced according to these criteria are enframed. **C**. Intracellular IFN-γ, TNF-α or IL-2 expression, representative examples (left panels) and summary of quantifications (right panels). Statistical significance was determined by ANOVA with Tukey’s post hoc test for panels **A** and **C**. Replicates are separately cultured and tested TIL products.

Because tumor reactivity was not detected in all TIL products (Fig. 4B), we asked whether the presence of tumor reactivity correlated with a particular *ex vivo* immune signature. Therefore, we compared the tumor immune cell composition to that of the kidney tissue (Fig. S5A). Even though the immune composition between digests greatly varied between patients, we detected significant differences in T cell, monocyte, neutrophil, DC and NKT cell content between tumor and kidney tissues (Fig. S5A). The most prominent of these differences was the reduced relative abundance of neutrophils in tumor digests (Fig. S5A). Because of the substantial variation and thus patient-specific immune landscape in RCC and kidney tissue, we calculated the difference between immune infiltrates in tumor and kidney digest and correlated that to the tumor reactivity of TIL products. Tumor reactivity was defined as follows: at least one of the individual TIL cultures from a patient showed >5% CD137 expression difference in CD8^+^ T cells comparing coculture with tumor digest versus medium control (ΔCD137hi) (Fig. 4B, blue line). TIL products with a ΔCD137^hi^ <5% were considered ‘not reactive’. We then correlated the reactive and non-reactive groups to changes in immune cell infiltration between tumor and kidney tissue. We found that tumor-reactive TIL products were more often derived from tumor tissues that contained increased infiltrates of T cells, NKT cells, B cells, and/or a loss of neutrophils (Fig. S5B). Of note, the percentages of these immune subsets just in tumor digests (Fig. 1B) lacked a significant correlation with tumor reactivity (Fig. S5C). Likewise, the percentage of tumor-infiltrating CD8^+^ T cells, CD4^+^ conv T cells, FOXP3^+^ CD25hi T cells (Fig. 1C), the percentage of PD-1hi T cells (Fig. 1D and E) and the tumor or immune cell PD-L1 expression (Table 1) did not correlate with tumor reactivity (Fig. S6A-C).

As the PCA of T cell infiltrates derived from tumor and kidney tissue showed key differences in the memory phenotype and the percentage of CD25 expressing CD8^+^ and CD4^+^ conv T cells (Fig. 2A-G), we also investigated whether these parameters correlated with the tumor reactivity of TIL products. The differentiation status of CD8^+^ and CD4^+^ conv T cells in tumor digests, whether or not in relation to kidney tissue, did not correlate with tumor reactivity of the in vitro expanded TIL product (Fig. S7A and B). The percentage of CD25 expressing T cells correlated with tumor reactivity of the TIL product only within the tumor-infiltrating CD8^+^ T cell population (Fig. S7C and D). Thus, increased T cell numbers that include CD25-expressing CD8^+^ T cells in the tumor tissue indicated an increased likelihood of generating tumor-reactive TIL products.

### Expanded tumor-reactive TILs fail to produce cytokines

Because effective TIL therapy requires cytokine-producing TIL products^29^, we examined whether expanded TILs also had the capacity to produce the key pro-inflammatory cytokines IFN-γ, TNF-α, and IL-2 when exposed to tumor tissue digest. Strikingly, even though PMA/ionomycin activation of TIL products showed the capacity of TIL products to produce high levels of IFN-γ, TNF-α, and IL-2 (Fig. 3D), and even though CD137 expression was clearly induced in TIL products from 10 out of 16 patients when exposed to autologous tumor digest (Fig. 4A and B), we failed to detect tumor-specific production of these three cytokines in both the CD8^+^ and the CD4^+^ T cells (Fig. 4C). In conclusion, even though tumor-reactive T cells in RCC lesions can be effectively expanded and produce cytokines after stimulation with pharmacological reagents, they fail to produce pro-inflammatory cytokines in response to tumor digests.

## Discussion

In this study, we show that TILs from RCC lesions can be effectively expanded with a standard REP. From 9 out of 16 patients (56%), TIL products contained tumor-reactive T cells as defined by the induction of tumor-specific CD137 expression. Our results are in the same range as seen in previous studies, although comparisons are complicated because of study-specific differences in methodology.^25–27^ The patient group we analyzed here contained RCC lesions from pretreated and treatment-naïve patients. While the study group is too small for firm conclusions, a clear relation between pretreatment and TIL expansion or tumor reactivity of the TIL product was absent.

Whether a tumor-reactive TIL product can be developed may depend on the composition of the tumor immune infiltrate. The comparison of the tumor immune landscape with that of non-tumor tissue from the same patient allowed us to identify tumor-specific infiltrations. Here we show that an enrichment of T cells in RCC compared to autologous kidney tissue positively correlated with the presence of tumor-reactive T cells in the TIL products. This is in apparent accordance with data in colon and ovarian tumors showing a positive association between high tumor T cell infiltration and clinical outcome.^37–39^ In addition, high expression of the activation markers PD-1, CD69 and CD25 on primary TILs is an important indicator for the percentage of tumor-reactive T cells and therapeutic response in NSCLC.^21,31^ In the current study, we only found that the presence of CD25-expressing CD8^+^ TILs correlated with tumor reactivity of expanded TILs.

Also, an increased presence of B cells in tumor lesions compared to paired kidney tissue associated with the occurrence of tumor-reactive T cells in TIL products. B cells are indicative for the presence of tertiary lymphoid structures, which are thought to drive anti-tumor immunity.^40^ In line with this, B cell signatures are enriched in responders to immune checkpoint blockade compared to non-responders in RCC, sarcoma and melanoma.^40,41^ Finally, a reduction of neutrophils in RCC lesions compared to paired kidney tissue correlated with increased tumor reactivity of expanded TIL products. Although only present in small numbers, it is tempting to speculate that neutrophils interfere with the functional potential of tumor-reactive T cells, especially because low neutrophil to lymphocyte ratios in RCC positively correlate with clinical anti-PD-1 therapy responses.^42^ Overall, we show that the composition of the tumor specific immune infiltrate in all likelihood determines the potential to generate a TIL product containing tumor-reactive T cells.

Although we found CD137-expressing tumor-reactive T cells, they generally failed to produce cytokines after stimulation with autologous RCC. Other pioneering studies reporting tumor reactivity of expanded RCC TILs showed low cytokine production and lacked thorough polyfunctional analyses.^25–27^ Differences between these and our studies are likely caused by the specific isolation and stimulation protocols. Intriguingly, with the identical protocol as we used here, we generated clinical-grade melanoma TILs at our facilities that readily produce effector cytokines against autologous tumor digest.^33^ Similarly, 76% of our NSCLC-derived TIL products showed tumor-specific cytokine responses to autologous tumors, of which 25% was even polyfunctional.^21^ Thus, the lack of tumor-specific cytokine production we observed for RCC-derived TIL products is tumor-type specific. The absence of tumor-specific cytokine production could not be attributed to an overall inability to produce these cytokines, as the RCC TILs produced ample amounts upon stimulation with PMA/ionomycin. Hence, what are the underlying mechanisms for the lack of cytokine production by RCC-derived TILs in response to tumor digests? A lack of cytokine production may have been imposed during the tumor reactivity assay by immune cells that include suppressive T cell populations or myeloid-derived suppressor cells (MDSCs). Even though we did not find a significant correlation of *ex vivo* CD4^+^FOXP3^+^CD25^hi^ cells with tumor reactivity, this T cell subset could be strongly suppressive.^43,44^ Alternatively, RCC-resident CD25-expressing CD4^+^FOXP3^-^ T cells can produce IL-10 and correlate with worse patient survival.^45^ In addition, abundantly present MDSCs in RCC negatively affect TIL outgrowth, and potentially interfere with immunotherapy.^28^ Furthermore, non-immune cells such as tumor cells or stromal cells often harbor the capacity to suppress functional T cell responses to tumor digest.^46^ For example, surface PD-L1 is one of the known key molecules to suppress anti-tumor recognition, and was expressed by tumor cells in some RCC lesions in this study. Alternatively, an immunosuppressive environment could hinder the re-programming of dysfunctional T cells to gain a pro-inflammatory profile. Because PMA/ionomycin activation induced strong responses, we consider this possibility unlikely. Nevertheless, it is conceivable that signaling nodules engaged upon TCR-HLA complex interaction that are required for cytokine production, but not for CD137 induction, may not have been completely rewired during the TIL cultures. Further investigations should reveal whether the generation of RCC-derived TIL products would benefit from the upfront removal of any or all of these immune suppressive cells.

In addition to the conclusions of other pioneering studies on opportunities of TIL therapy for the treatment of RCC,^25–27^ we recommend that TIL generation from RCC first requires a thorough mechanistic understanding of the RCC TIL dysfunction before application to patients. Moreover, the lack of functionality of RCC TILs may have contributed to the failure of their clinical application in the late 90s.^47^ As of writing, we could only find one active TIL clinical trial for RCC in the ClinicalTrials.gov database.^48^ In contrast, multiple checkpoint inhibitor trials are ongoing, further emphasizing the current relative lack of knowledge to design effective TIL therapy for RCC.^49–51^

In conclusion, we successfully expanded tumor-reactive TILs from RCC patients, but further elucidation of molecular and cellular suppressive pathways in RCC is needed to restore their tumor-specific cytokine production.

## Acknowledgments

We thank the flow cytometry facility of Sanquin Research, and the medical assistance staff from The Netherlands Cancer Institute – Antoni van Leeuwenhoek hospital (Amsterdam) for technical help. This work was supported by intramural funding of Stichting Sanquin Bloedvoorziening (PPOC 14-46).

The authors declare no potential conflicts of interest.

