## Supplementary Figures and Legends for "T cells expanded from renal cell carcinoma are tumor-reactive but fail to produce IFN-γ, TNF-α or IL-2"


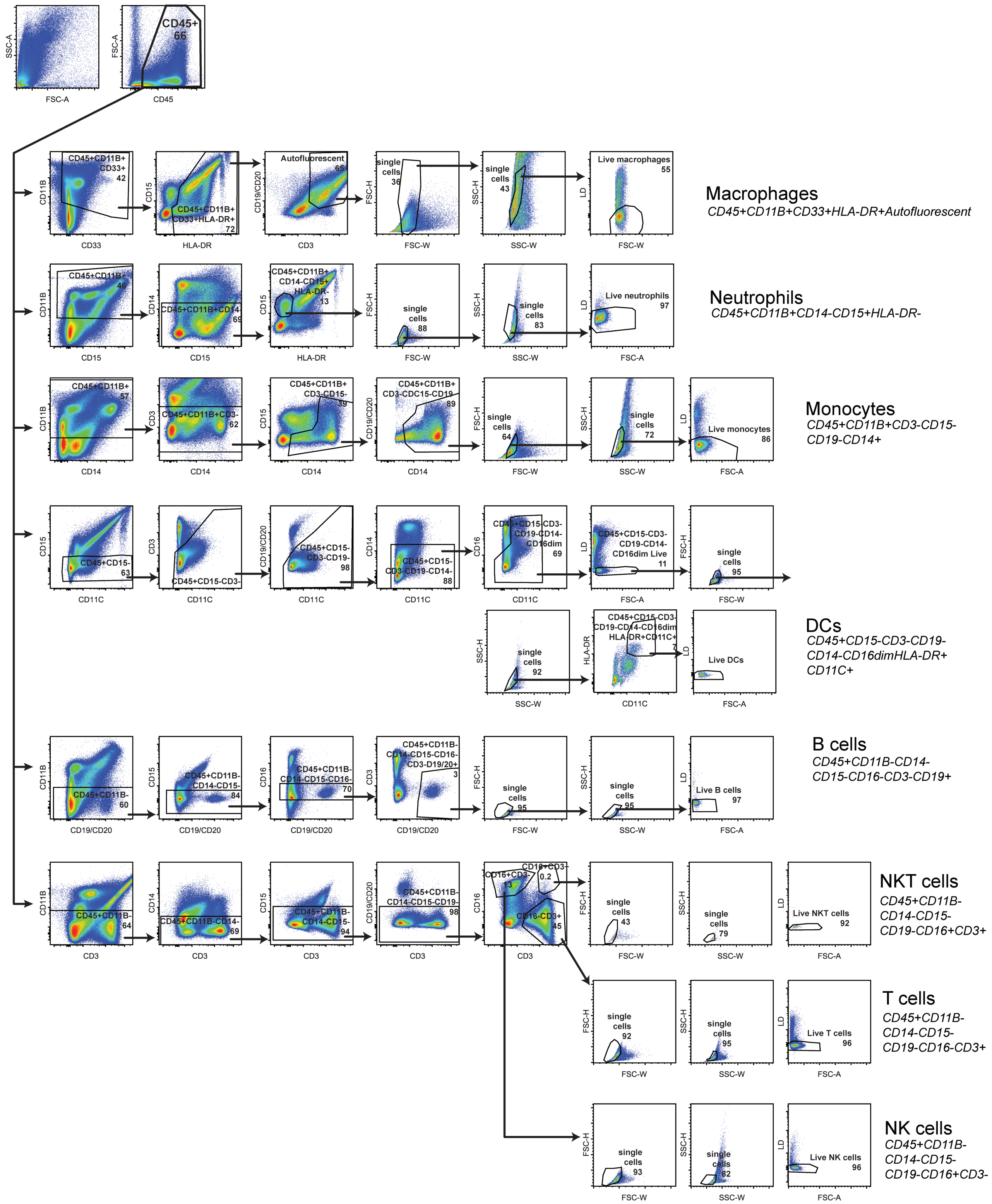


**Fig. S1 Gating strategy immune cell subsets.** Gating strategy to determine the major immune subsets as quantified in Fig. 1. All subsets were first gated positively for CD45. Macrophages were gated positively for CD11b, HLA-DR, autofluorescence, followed by doublet exclusion and dead cell exclusion. Neutrophils were gated positively for CD11b, negatively for CD14, $CD15^{+}HLA\text{-}DR^{-}$, followed by doublet exclusion and dead cell exclusion. Monocytes were gated positively for CD11b, negatively for CD3, $CD15^{+}CD14^{-}$ cells were excluded, then gated positively for CD14, followed by exclusion of doublets and dead cells. Dendritic cells (DCs) were gated negatively for CD15, CD3, CD19/CD20 and CD14. $CD16^{+}CD11C^{-}$ cells, dead cells and doublets were excluded. Finally, DCs were identified as $CD11C^{+}HLA\text{-}DR^{+}$. B cells were gated negatively for CD11b, CD15, CD16, and positively for CD19/CD20. Doublets and dead cells were excluded. T cells and NK(T) cells had a similar gating strategy. They were gated negatively for CD11B, CD14, CD15 and CD19/CD20. Next NK cells were identified as $CD3^{-}CD16^{+}$, NKT cells as $CD3^{+}CD16^{+}$ and T cells as $CD3^{+}CD16^{-}$. Doublets and dead cells were excluded.

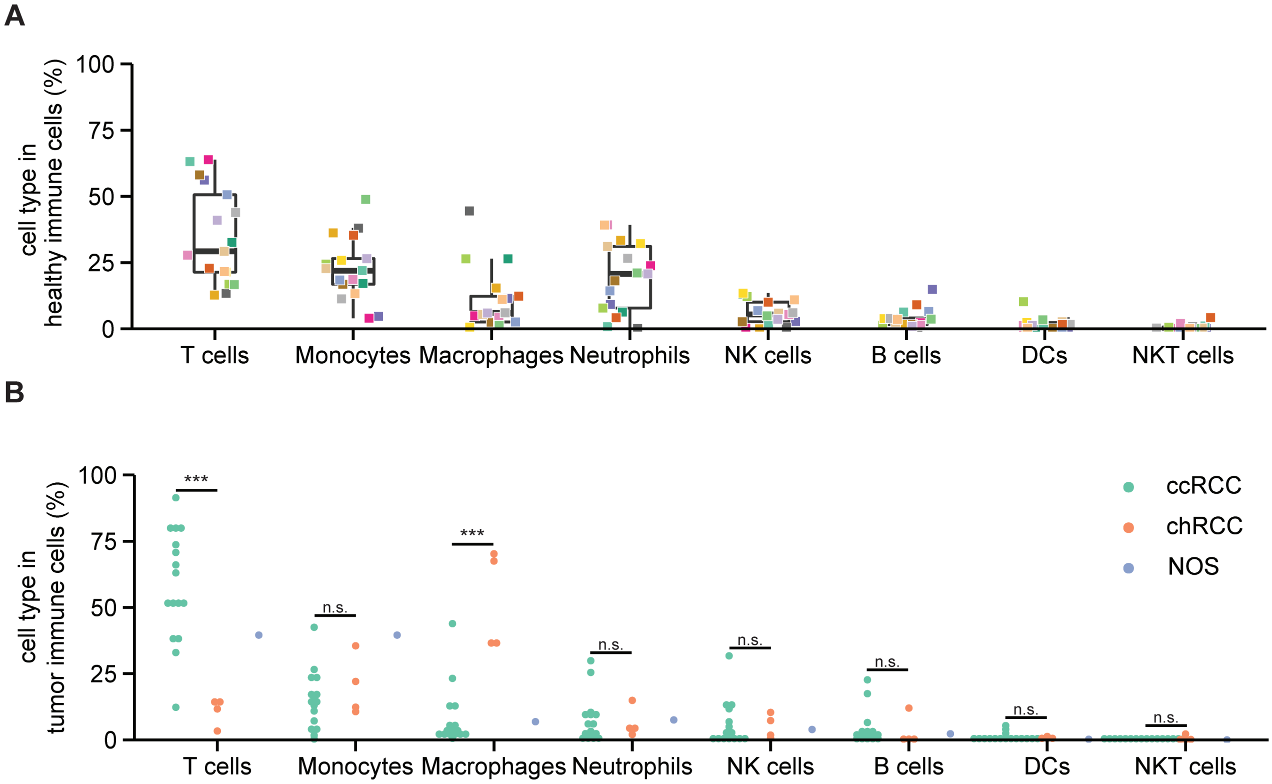


**Fig. S2 Immune subsets in kidney and in tumor per RCC subtype**. **A-B**. Immune subsets as a percentage of total immune cells in kidney digest (**A**) and in tumor digest per RCC type (**B**). ccRCC = clear cell RCC, chRCC = chromophobe RCC, NOS = not otherwise specified. Aligned ranks transformation two-way ANOVA, Tukey's post hoc test.

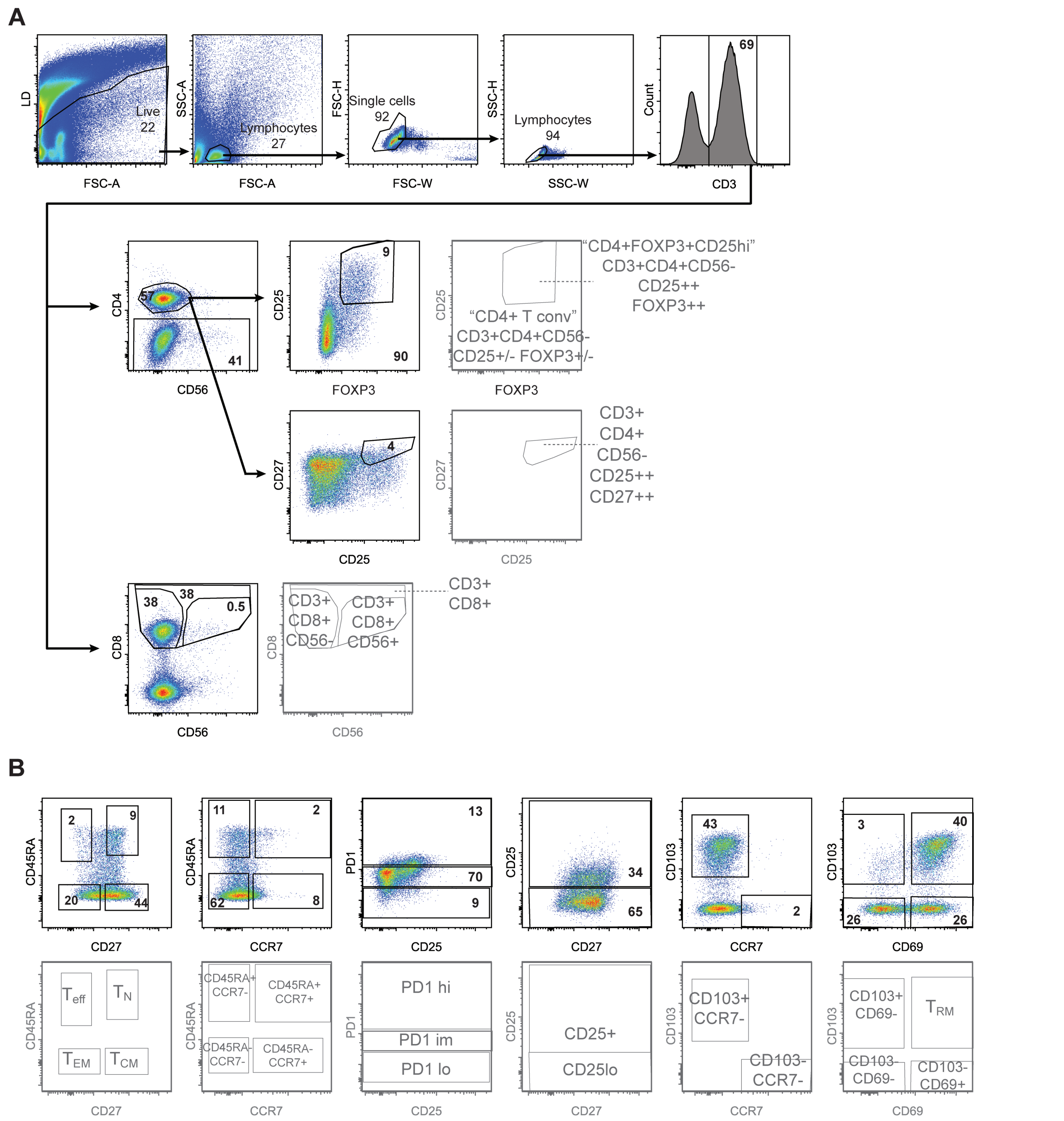


**Fig. S3. Gating strategy of the T cell focused antibody panel. A.** Lymphocytes were first gated negatively for the life/dead stain, then gated based on cell size, followed by exclusion of doublets. T cells were identified as CD3$^+$. Within the CD8$^+$ T cell population, CD8$^+$CD56$^-$ and CD8$^+$CD56$^+$ subpopulations were identified. The CD4$^+$ T cell population was split into FOXP3$^+$CD25$^{hi}$ and non-FOXP3$^+$C-D25$^{hi}$ (CD4$^+$ conv) subpopulations. **B**. Gating strategy for the determination of various memory and activation phenotypes. This gating strategy was applied to all populations defined in A. This antibody panel was used in Figs. 1C-D, 2, S6 and S7.

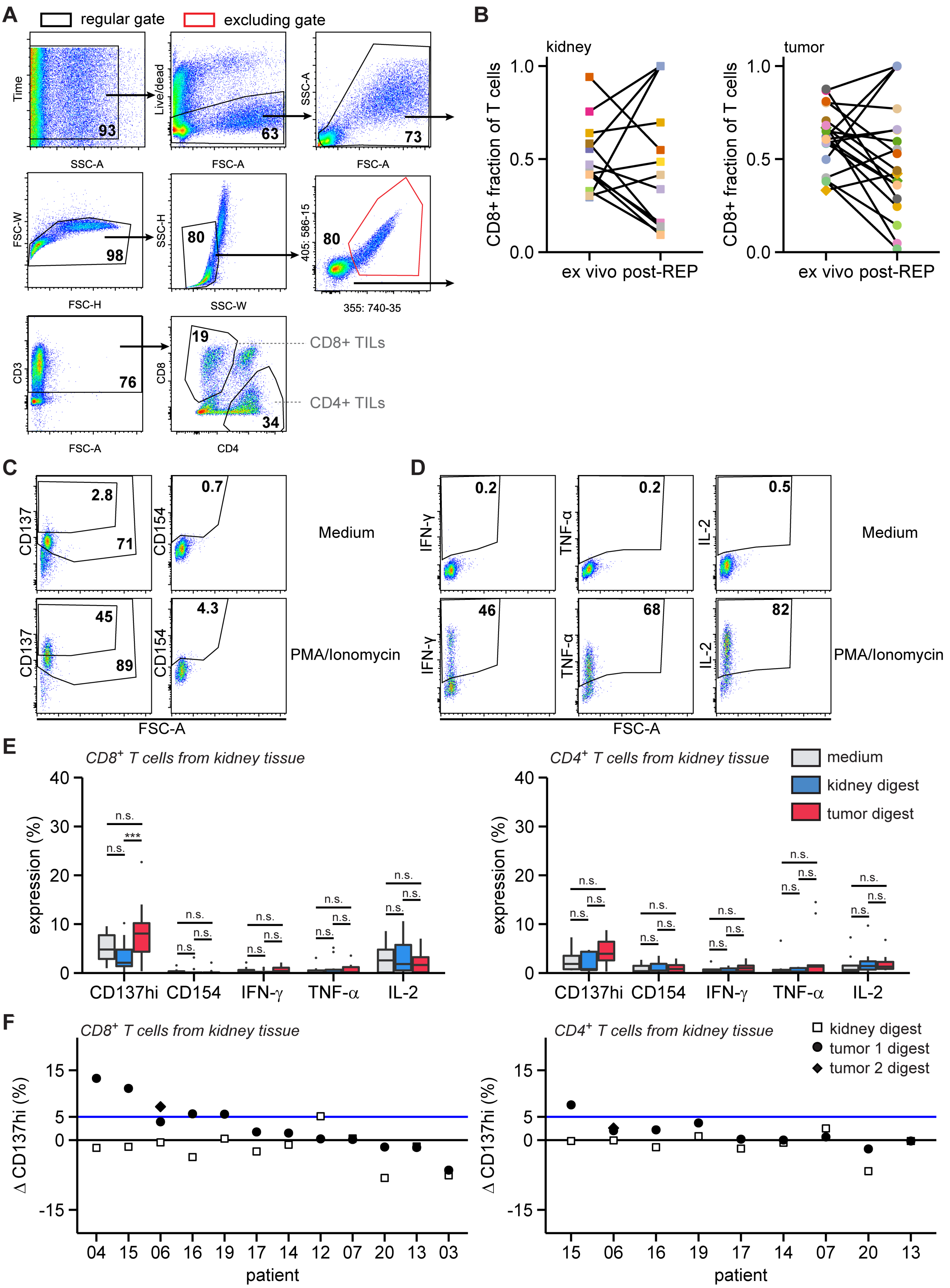


**Fig. S4. Gating strategy tumor reactivity assay and tumor reactivity TIL products from kidney. A.** Any obstructions in the flow of the cytometer were gated out using a time gate. Next, dead cells were excluded, cells were gated based on size followed by doublet exclusion. Tumor and kidney digests often contained autofluorescent cells. These cells were removed by excluding cells positive in two empty channels. Finally, TILs were identified by expression of CD3, and CD4 or CD8. **B.** Fraction of CD8⁺ T cells within total T cells per patient (corresponds to Fig. 3B). **C.** Example gating strategies for the activation markers CD137 and CD154. **D.** Example gating strategies for the cytokines IFN-γ, TNF-α or IL-2. **E-F.** TIL products derived from kidney digests were cultured in medium with or without autologous tumor or kidney digest for 6 h. Extracellular CD154 and intracellular high CD137 (CD137hi), IFN-γ, TNF-α or IL-2 expression per stimulation (**E**) and per patient delta CD137hi (Δ; tissue digest – medium) reactivity of TIL products (**F**).

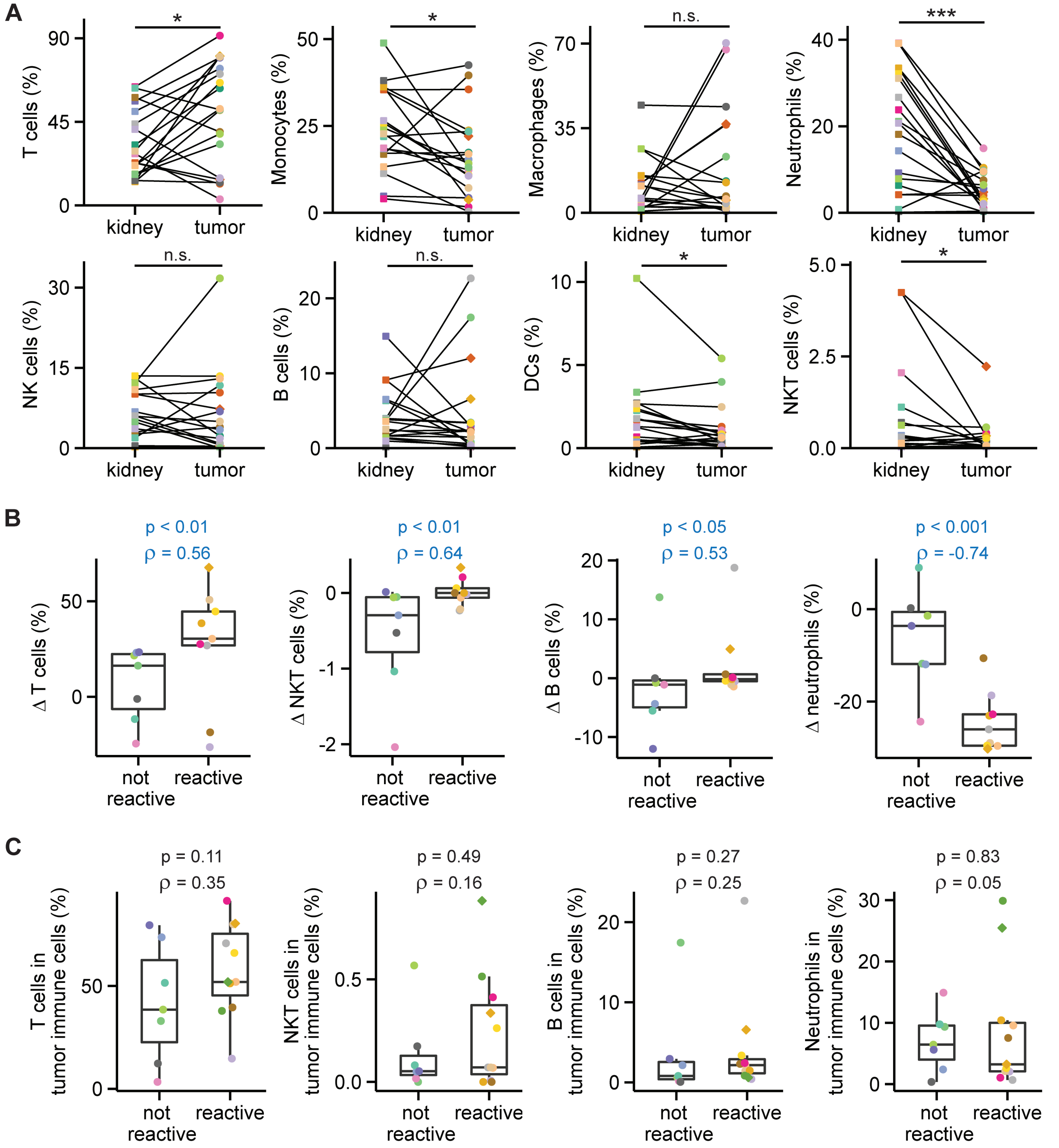


**Fig S5. T cell infiltration correlates with tumor reactivity. A.** *Ex vivo* immune subsets as a percentage of total immune cells for paired kidney and tumor samples (data from Fig. 1B and S2A). Statistical significance was determined by Wilcoxon signed-rank test. **B.** Delta (**Δ**) *ex vivo* immune cell subsets (tumor – kidney) were correlated with post-REP tumor reactivity of TIL products (Fig. 4B) using Spearman's correlation. **C.** Spearman's correlation between the percentage of immune subsets in the *ex vivo* RCC samples (Fig. 1B) and post-REP tumor reactivity (Fig. 4B). For **C** and **D** p values and Spearman's rho (ρ) are shown above each panel. Significant ($p < 0.05$) correlations are colored blue, nonsignificant correlations black.

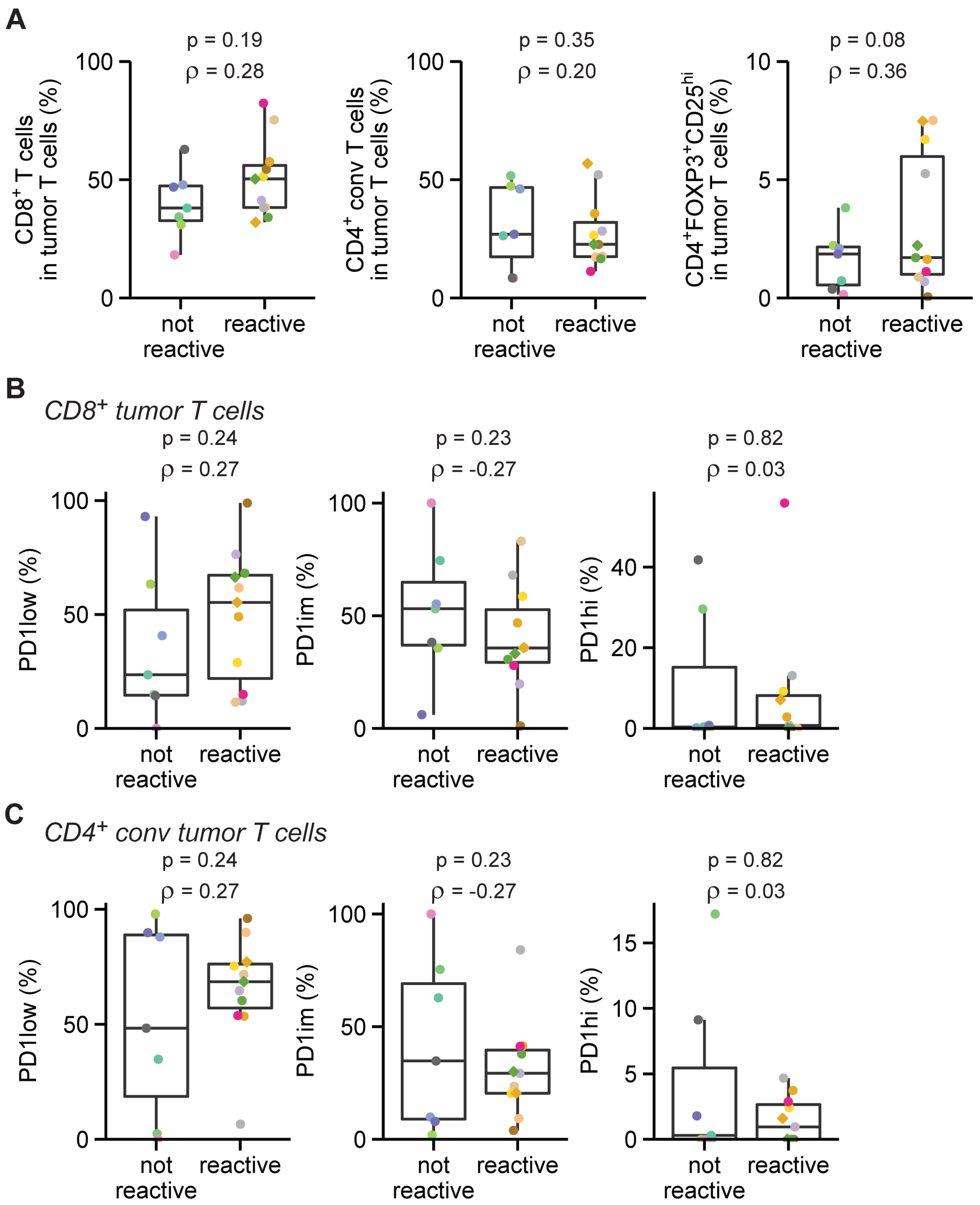


**Fig S6. T cell subsets nor T cell PD-1 expression correlate with tumor reactivity. A.** Spearman's correlation between *ex vivo* T cell subpopulations (Fig. 1C) and post-REP tumor reactivity of TIL products (Fig. 4B). **B-C** Spearman's correlation between PD-1 expression levels in $CD8^+$ T cells (Fig. 1D) (**B**) or $CD4^+$ T cells (Fig. 1E) (**C**) and post-REP tumor reactivity (Fig. 4B). P values and Spearman's rho (ρ) are shown above each panel. Significant correlations ($p < 0.05$) are colored blue, nonsignificant correlations black.

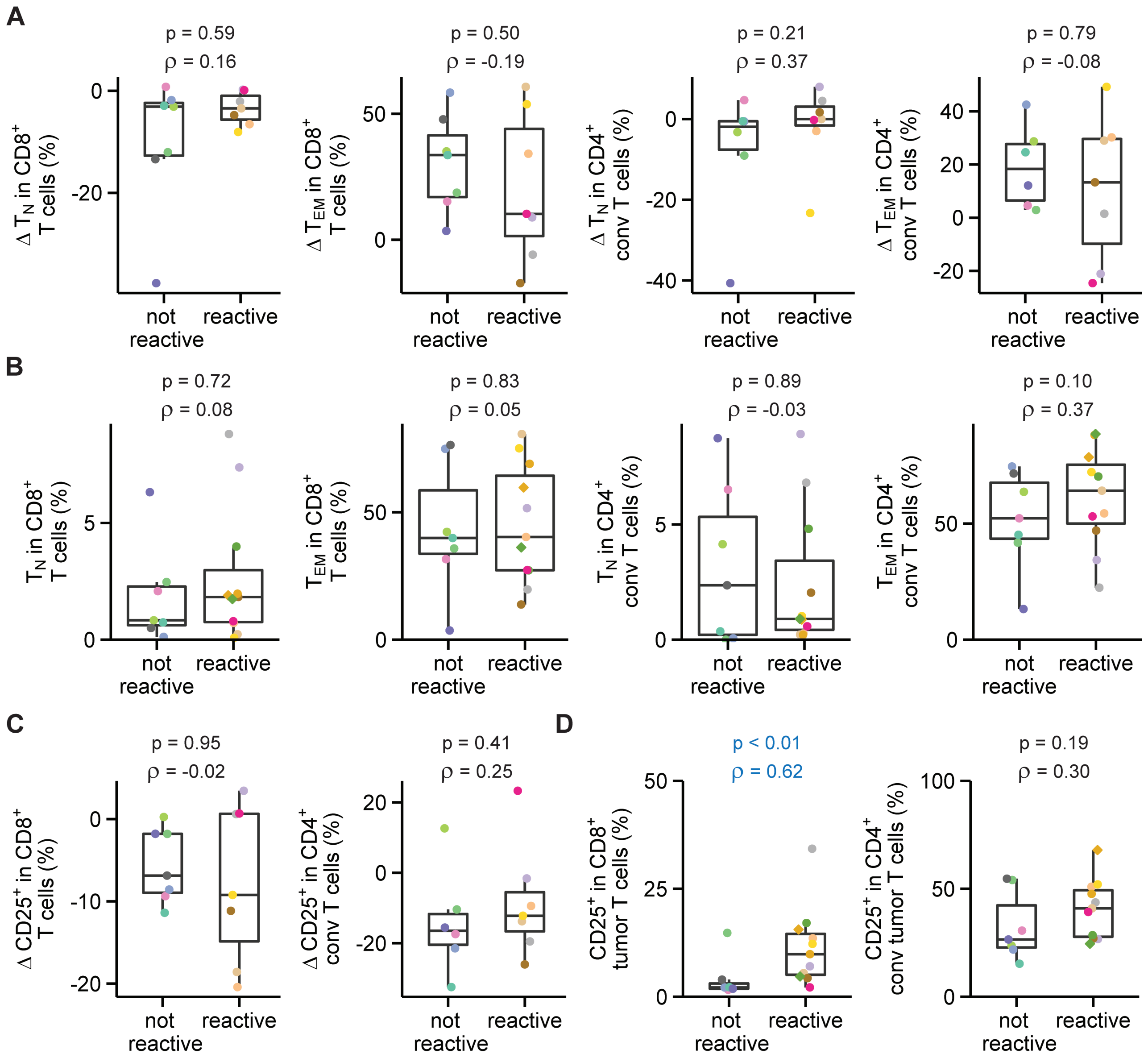


**Fig S7. CD25 expression in ex vivo tumor CD8+ T cells correlates with tumor reactivity. A-B.** *Ex vivo* naïve ($CD45RA^+CD27^+$) and effector memory ($CD45RA^-CD27^-$) populations within $CD8^+$ and $CD4^+$ conv T cells (Fig. 2D and E) were correlated with post-REP tumor reactivity of TIL products (Fig. 4B) using Spearman's correlation. Correlation of the difference in infiltration between tumor and kidney **(**Δ; tumor - kidney) and tumor reactivity is shown in **A**, the correlation between tumor T cell subsets and tumor reactivity in **B**. **C-D.** *Ex vivo* CD25 expression in $CD8^+$ and $CD4^+$ conv T cells (Fig. 2F and G) correlated with post-REP tumor reactivity (Fig. 4B). Delta CD25 expression (tumor – kidney) is shown in **C**, tumor CD25 expression in **D**. P values and Spearman's rho (ρ) are shown above each panel. Significant correlations ($p < 0.05$) are colored blue, nonsignificant correlations black.
